# Microbial biogeography of 1,000 geothermal springs in New Zealand

**DOI:** 10.1101/247759

**Authors:** J.F. Power, C.R. Carere, C.K. Lee, G.L.J. Wakerley, D.W. Evans, M. Button, D. White, M.D. Climo, A.M. Hinze, X.C. Morgan, I.R. McDonald, S.C. Cary, M.B. Stott

## Abstract

Geothermal springs are model ecosystems to systematically investigate microbial biogeography as they i) represent discrete, homogenous habitats; ii) are abundantly distributed across multiple geographical scales; iii) span broad geochemical gradients; and iv) have simple community structures with reduced metazoan interactions. Taking advantage of these traits, we undertook the largest known consolidated study of geothermal ecosystems (http://1000springs.org.nz) to determine factors that influence biogeographical patterns. Rigorously standardised methodologies were used to measure microbial communities, 46 physicochemical parameters, and metadata from 1,019 hotspring samples across New Zealand. pH was found to be the primary influence on diversity in springs < 70 °C with community similarity decreasing with geographic distance. Surprisingly, community composition was dominated by two genera (*Venenivibrio* and *Acidithiobacillus*) in both average relative abundance (11.2 and 11.1 %) and prevalence (74.2 and 62.9 % respectively) across physicochemical spectrums of 13.9 – 100.6 °C and pH < 1 – 9.7. This study provides an unprecedented insight into the ecological conditions that drive community assembly in geothermal springs, and can be used as a foundation to improve the characterisation of global microbial biogeographical processes.

---

Microbial biogeography identifies patterns of diversity across defined spatial or temporal scales in an attempt to describe the factors which influence these distributions. The pervasive view that microorganisms are dispersed ubiquitously and therefore do not adhere to classical biogeographical patterns has been historically presumed^1^. Recent studies, however, have contradicted this paradigm and shown that microbial community diversity is shaped across time and space^2,3^ via a combination of environmental selection, stochastic drift, diversification and dispersal limitation^4,5^. The relative impact of these ecological drivers on diversity is the subject of ongoing debate, with differential findings reported across terrestrial, marine and human ecosystems^6-12^.

Geothermally-heated springs are ideal systems to investigate microbial biogeography. In comparison to terrestrial environments, geothermal springs represent discrete, homogenous aquatic habitats with broad physicochemical gradients distributed across proximal and distal geographic distances. The relatively simple microbial community structures, typical of geothermal springs, also allow for the robust identification of diversity trends. Separate studies have each alternatively implicated temperature^8,13,14^, pH^15^, and seasonality^16^ as the primary drivers of community diversity in these ecosystems; with niche specialisation observed within both local and regional populations^17,18^. The neutral action of microbial dispersal is also thought to be a significant driver behind the distribution of microorganisms^23^, with endemism and allopatric speciation reported in intercontinental hotsprings^21,22^. It is important to note that significant community differences have been found between aqueous and soil/sediment samples from the same springs^13,15,23^, emphasising that the increased relative homogeneity of aqueous samples make geothermal water columns excellent candidate environments for investigating large scale taxa-geochemical associations. However, despite these findings, a lack of scale (e.g.geographic distance, sampling quantity/density and physicochemical gradients) and uniformity in sampling methodology has hindered a holistic view of microbial biogeography from developing.

The Taupō Volcanic Zone (TVZ) is a region rich in geothermal hotsprings and broad physicochemical gradients spanning 8,000 km^2^ in New Zealand’s North Island (Fig. 1), making it a tractable model system for studying microbial biogeography. This unique area is a rifting arc associated with subduction at the Pacific-Australian tectonic plate boundary, resulting in a locus of intense magmatism^24^. The variable combination of thick, permeable volcanic deposits, high heat flux, and an active extensional (crustal thinning) setting favours the deep convection of groundwater and exsolved magmatic volatiles that are expressed as physicochemically-heterogeneous surface features in 23 geographically distinct geothermal fields^25,26^. Previous microbiological studies across the region have hinted at novel diversity and function present within some of these features^27-31^, however investigations into the biogeographical drivers within the TVZ are sparse and have focused predominantly on soil/sediments or individual hotsprings^8,14,32^.

**Fig. 1.**
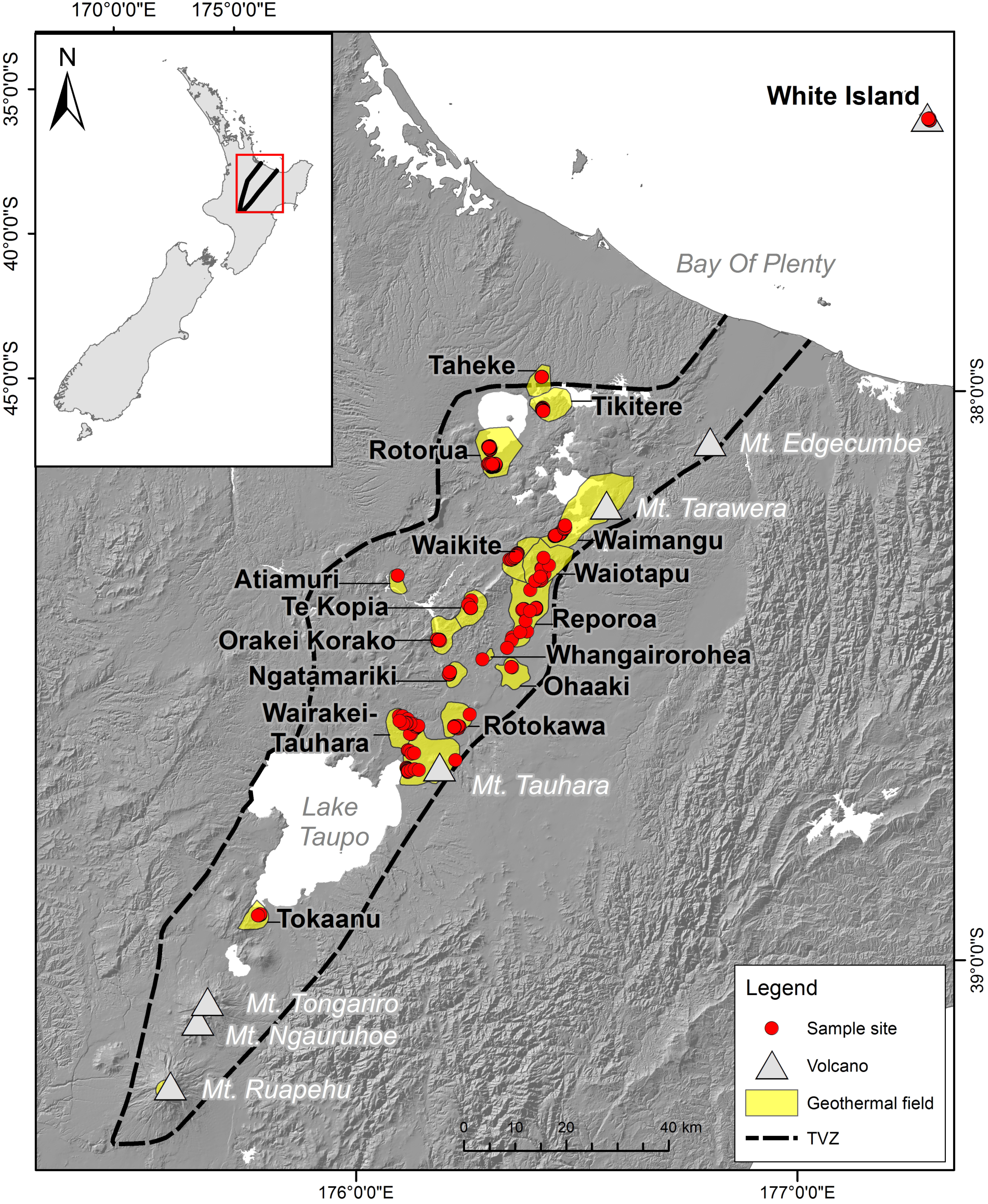
Map of the Taupō Volcanic Zone (TVZ), New Zealand. Geothermal fields are highlighted in yellow, with springs sampled for the 1,000 Springs Project in red (*n*= 1,019).

Here we report the diversity and biogeography of microbial communities found in over 1,000 geothermal spring samples, collected as part of the 1,000 Springs Project. This project aimed to catalogue the microbial biodiversity and physicochemistry of New Zealand’s iconic hotsprings to serve as a conservation, scientific, and indigenous cultural knowledge repository for these ecosystems. A publicly accessible database of all springs surveyed is available online (https://1000Springs.org.nz). Over a period of 93 weeks, rigorously standardised methodologies were used to collect samples/metadata, perform community analysis and quantify physicochemistry within the TVZ to answer the following three questions:

1. To what extent does physicochemistry and geography influence microbial diversity and community structure within geothermal springs?
2. How does the influence of significant physicochemical parameters change in response to the gradation of other major community drivers?
3. Can taxon-specific geochemical niches be identified for abundant microorganisms in these ecosystems?

This work represents the largest known microbial ecology study on geothermal aquatic habitats at a regional scale. Our results clearly demonstrate both the relative influence of physicochemical parameters (e.g. pH) and the effect of geographic isolation on the assemblage of communities in these extreme ecosystems. Collectively these findings expand our knowledge of the constraints that govern universal microbial biogeographical processes.

## Results & discussion

Recent biogeography research has demonstrated microbial diversity patterns are detectable and are influenced by both deterministic^33^ and stochastic processes^6^. A lack of consensus on the relative impact of these factors, however, has been exacerbated by an absence of broad physicochemical gradients, and sampling scale and density across both geographic distance and habitat type. The inherent heterogeneity of terrestrial soil microbial ecosystems^34,35^ further confounds attempts to distinguish true taxa-geochemical associations. To provide greater resolution to the factors driving microbial biogeography processes, we determined the physicochemical and microbial community composition of 1,019 geothermal water-column samples from across the TVZ (Fig. 1). Samples included representatives of both extreme pH (< 0 – 9.7) and temperature (13.9 – 100.6 °C) (Supplementary Fig. 1). The filtering of low-quality and temporal samples yielded a final data set of 925 individual geothermal springs for spatial-statistical analysis (more details can be found in the Supplementary Methodology). From these 925 springs, a total of 28,381 operational taxonomic units (OTUs) were generated for diversity studies.

### Microbial diversity is principally driven by pH, not temperature, in geothermal spring ecosystems

Reduced microbial diversity in geothermal springs is often attributed to the extreme environmental conditions common to these areas. Temperature and pH are reported to be the predominant drivers of microbial diversity^8,36^, but their influence relative to other parameters has not been investigated over large geographic and physicochemical scales with appropriate sample density. Our analysis of microbial richness and diversity showed significant variation spanning pH, temperature and geographical gradients within the TVZ (richness: 49 – 2997 OTUs, diversity: 1.1 – 7.3 Shannon index; Supplementary Fig. 2&3). As anticipated, average OTU richness (386 OTUs; Supplementary Fig. 4) was substantially reduced in comparison to studies of non-geothermal temperate terrestrial^37,38^ and aquatic^39^ environments. Further, OTU richness was maximal at the geothermally-moderate temperature of 21.5 °C and at circumneutral pH 6.4. This is consistent with the hypothesis that polyextreme habitats prohibit the growth of most microbial taxa, a trend reported in both geothermal and non-geothermal environments alike^8,12^. A comparison of linear regressions of 46 individual physicochemical parameters (Supplementary Table 1) confirmed pH as the most significant factor influencing diversity (16.4 %, *n* = 925, *p*< 0.001; Supplementary Fig. 3), while further multiple regression analysis showed 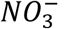, turbidity, oxidation-reduction potential (ORP), dissolved oxygen, 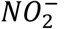, *Si* and *Cd* also had meaningful contributions (Supplementary Table 2). Cumulatively, along with pH, these factors accounted for 26.6 % of the observed variation in Shannon diversity. Correlation of pH with Shannon index (Pearson’s coefficient: |*r*| = 0.41, *P* < 0.001) and significance testing between samples binned by pH increments (Kruskal-Wallis: *H* = 179.4, *P* < 0.001) further confirmed pH as a major driver of variation in alpha diversity. This finding is consistent with reports of pH as the primary environmental predictor of microbial diversity in several ecosystems (e.g. soil^12^, freshwater^40^, alpine^38^).

It has been previously hypothesised that pH has significant influence on microbial community composition because changes in proton gradients will drastically alter nutrient availability, metal solubility, or organic carbon characteristics^12^. Similarly, acidic pH will also reduce the number of taxa observed due to the low number that can physiologically tolerate these conditions. Here, we demonstrate that pH had the most significant effect on diversity across all springs measured, but due to our high sampling frequency, we see this influence reduced above 70 °C (Fig. 2). Inversely, the effect of temperature on diversity was diminished in springs where pH was <4 (Supplementary Fig. 5). There is some evidence that suggests thermophily predates acid tolerance^41,42^, thus it is possible the added stress of an extreme proton gradient across cell membranes has constrained the diversification of the thermophilic chemolithoautotrophic organisms common to these areas^43^. Indeed, a recent investigation of thermoacidophily in archaea suggests hyperacidophily (growth < pH 3.0) may have only arisen as little as ~ 0.8 *Ga*^42^, thereby limiting the opportunity for microbial diversification; an observation highlighted by the paucity of these microorganisms in extremely acidic geothermal ecosystems^14,42^. It is also important to note that salinity has previously been suggested as an important driver of microbial community diversity^44,45^. The quantitative data in this study showed only minimal influence of salinity (proxy as conductivity) on diversity (Supplementary Table 1), bearing in mind that the majority of the hotspring samples in this study had salinities substantially less than that of seawater.

**Fig. 2.**
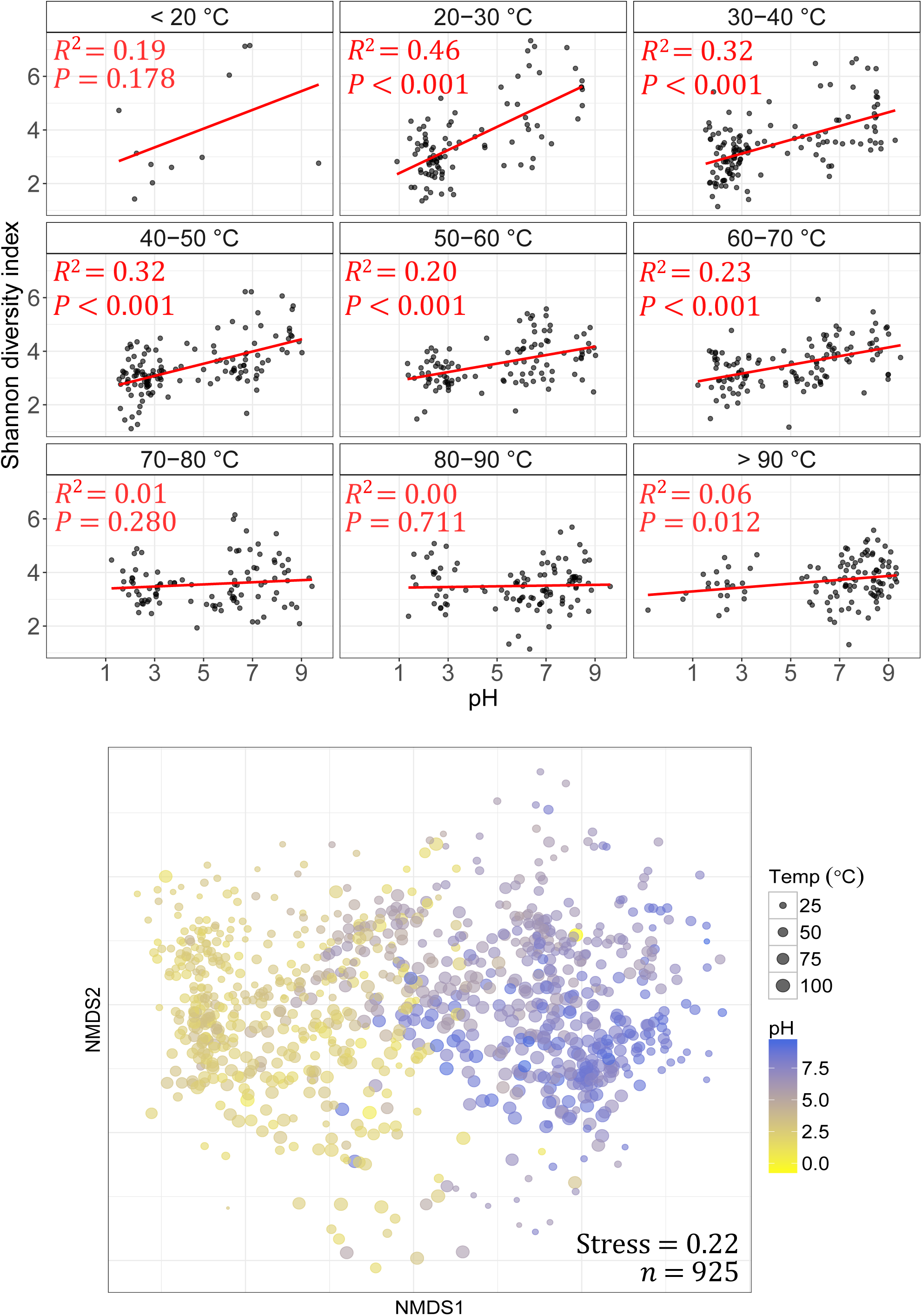
Alpha and beta diversity against pH and temperature. (Top) pH against alpha diversity via Shannon index of all individual springs (*n*= 925) in 10 °C increments, with linear regression applied to each increment. (Bottom) Non-metric multidimensional scaling (NMDS) plot of beta diversity (via Bray-Curtis dissimilarities) between all microbial community structures.

The relationship between temperature and diversity reported in this research starkly contrasts a previous intercontinental study comparing microbial community diversity in soil/sediments from 165 geothermal springs^8^, which showed a strong relationship (*R*^2^ = 0.40–0.44) existed. In contrast, our data across the entire suite of samples, revealed temperature had no significant influence on observed community diversity (*R*^2^ = 0.002, *P*= 0.201; Supplementary Fig. 3, Supplementary Table 1). This result increased marginally for archaeal-only diversity (*R*^2^ = 0.013, *P* = 0.0005), suggesting temperature has a more profound effect on this domain than bacteria. However, the primers used in this study are known to be unfavourable towards some archaeal clades^46^, therefore it is likely extensive archaeal diversity remains undetected in this study. The lack of influence of temperature on whole community diversity was further substantiated via multiple linear modelling (Supplementary Table 2), and significance and correlation testing (Kruskal-Wallis: *H* = 16.2, *P* = 0.039; Pearson’s coefficient: |*r*| = 0.04, *P* = 0.201). When samples were split into pH increments, like Sharp *et al.* (2014)^8^, we observed increasing temperature only significantly constrained diversity above moderately acidic conditions (pH > 4; Supplementary Fig. 5). However, the magnitude of this effect was, in general, far less than previously reported and is likely a consequence of the sample type (e.g. soil/sediments versus aqueous) and density processed^15^. Many geothermal environments are recalcitrant to traditional DNA extraction protocols and research in these areas has therefore focused on samples with higher biomass abundance^8,36^ (i.e. soils, sediments, streamers or biomats). Whereas aqueous samples typically exhibit a more homogenous chemistry and community structure, the heterogeneity of terrestrial samples is known to affect microbial populations (e.g. particle size, depth, nutrient composition)^34,35,47^. Our deliberate use of aqueous samples extends the results of previous small-scale work^13,32^ and also permits the robust identification of genuine taxa-geochemical relationships in these environments.

### Community structures are influenced by pH, temperature and geothermal source fluid

Throughout the TVZ, beta diversity correlated more strongly with pH (Mantel: *ρ* = 0.54, P< 0.001) than with temperature (Mantel: *ρ* = 0.19, *P* < 0.001; Fig. 2, Supplementary Table 3). This trend was consistent in pH- and temperature-binned samples (Supplementary Fig. 7; ANOSIM: |*R*| = 0.46 and 0.18 respectively, *P* < 0.001); further confirming pH, more so than temperature, accounted for observed variations in beta diversity. Congruent with our finding that pH influences alpha diversity at lower temperatures (< 70 °C), the effect of temperature reducing beta diversity had greater significance above 80 °C (*P* < 0.001; Supplementary Fig. 7). The extent of measured physicochemical properties across 925 individual habitats, however, allowed us to explore the environmental impact on community structures beyond just pH and temperature. Permutational multivariate analysis of variance in spring community assemblages showed that pH (12.4 %) and temperature (3.9 %) had the greatest contribution towards beta diversity, followed by ORP (1.4 %), 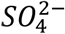 (0.8 %), turbidity (0.8 %) and *As* (0.7 %) (*P*< 0.001; Supplementary Table 4). Interestingly, constrained correspondence analysis of the 15 most significant, non-collinear and variable parameters (Supplementary Table 4 & 5; pH, temperature, turbidity, ORP, 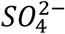, 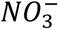, *As*, 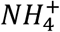, 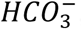, *H_2_S*, conductivity, *Li*, *Al*, *Si* and 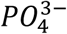), along with geothermal field locations, only explained 10 % of variation in beta diversity (Fig. 3), indicating physicochemistry, or at least the 46 parameters measured were not the sole drivers of community composition.

**Fig. 3.**
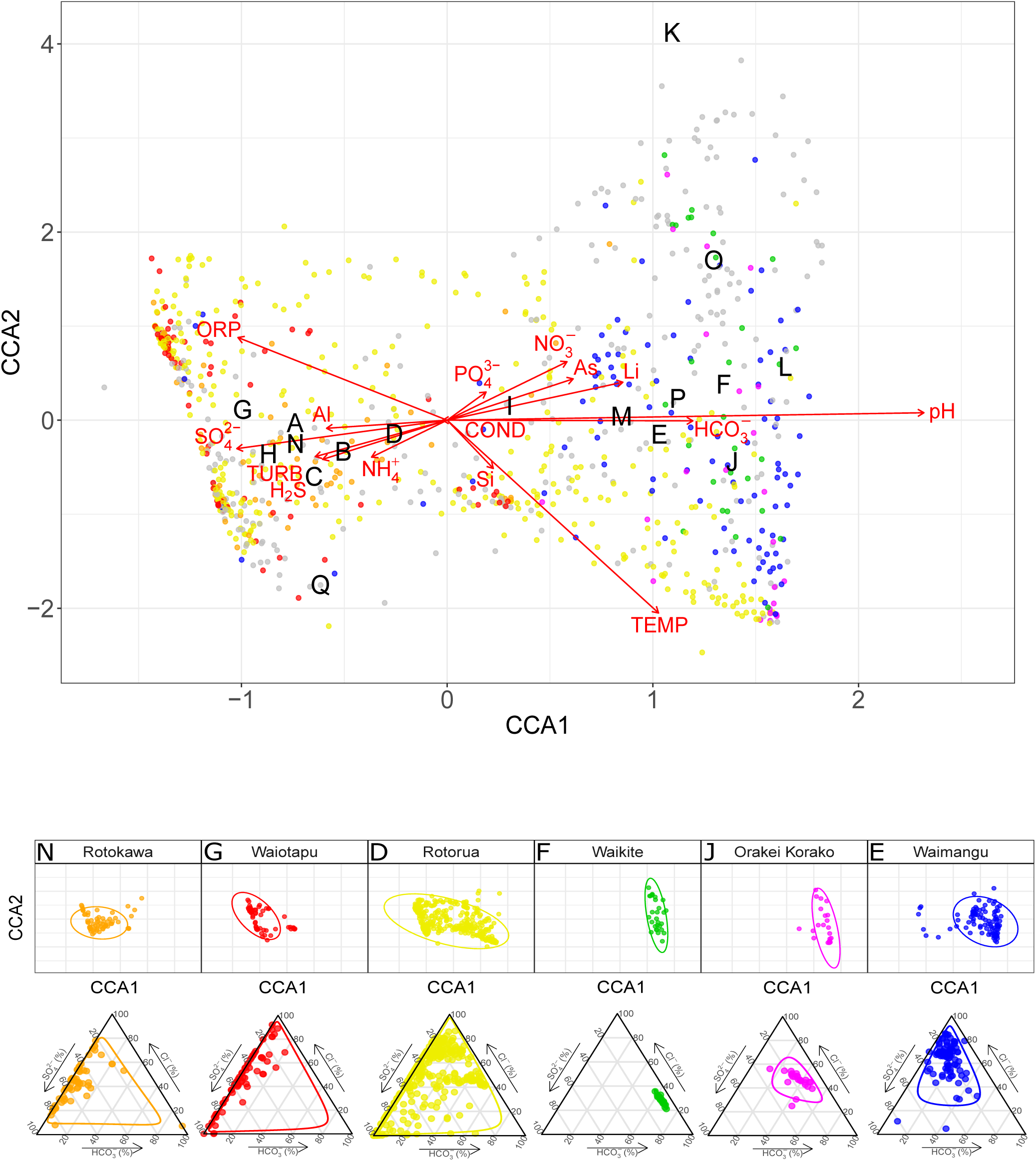
Constrained correspondence analysis (CCA) of beta diversity with significant physicochemistry. (Top) A scatter plot of spring community dissimilarities (*n* = 923), with letters corresponding to centroids from the model for geothermal fields (A-Q; White Island, Taheke, Tikitere, Rotorua, Waimangu, Waikite, Waiotapu, Te Kopia, Reporoa, Orakei Korako, Whangairorohea, Ohaaki, Ngatamariki, Rotokawa, Wairakei-Tauhara, Tokaanu, Misc). Coloured communities are from fields represented in the subpanel. Constraining variables are plotted as arrows (COND: conductivity, TURB: turbidity), with length and direction indicating scale and area of influence each variable had on the model. (Bottom) A subset of the full CCA model, with select geothermal fields shown in colour (including 95 % confidence intervals) and their respective geochemical signature as a ratio of chloride (Cl ^-^), sulfate 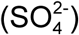 and bicarbonate 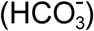.

We also investigated whether typical geochemical conditions exist for springs within the same geothermal field and whether specific microbial community assemblages could be predicted. Geothermal fields are known to express chemical signatures characteristic of their respective source fluids^48^, implying autocorrelation could occur between location and geochemistry. Springs are usually classified according to these fluids; alkaline-chloride or acid-sulfate. High-chloride features are typically sourced from magmatic waters and have little interaction with groundwater aquifers. At depth, water-rock interactions can result in elevated bicarbonate concentrations and, consequently, neutral to alkaline pH in surface features. Acid-sulfate springs (pH 2-3), in contrast, form as steam-heated groundwater couples with the eventual oxidation of hydrogen sulfide into sulfate (and protons). Rarely, a combination of the two processes can occur; leading to intermediate pH values^49^. It is unknown, however, whether these source fluid characteristics are predictive of their associated microbial ecosystems. Bray-Curtis dissimilarities confirmed that, like alpha diversity (Kruskal-Wallis: *H* = 240.7, *P* < 0.001; Fig. 5), community structures were significantly different between geothermal fields (ANOSIM: | *R* | = 0.26, *P* < 0.001; Supplementary Fig. 6). Gradient analysis comparing significant geochemical variables and geography further identified meaningful intra-geothermal field clustering of microbial communities (95 % *CI*; Fig. 3 & Supplementary Fig. 9). Further, characteristic geochemical signatures from these fields were identified and analysis suggests they could be predictive of community composition. For example, the Rotokawa and Waikite geothermal fields (approx. 35 km apart) (Fig. 3N & 3F) display opposing ratios of 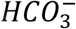, 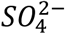 and *Cl^−^*, with corresponding microbial communities for these sites clustering independently in ordination space. Despite this association, intra-field variation in both alpha and beta diversity also occurred at other geothermal sites where geochemical signatures were not uniform across local springs (e.g. Rotorua, Fig. 3D), demonstrating that correlation does not necessarily always occur between locational proximity and physicochemistry.

**Fig. 4.**
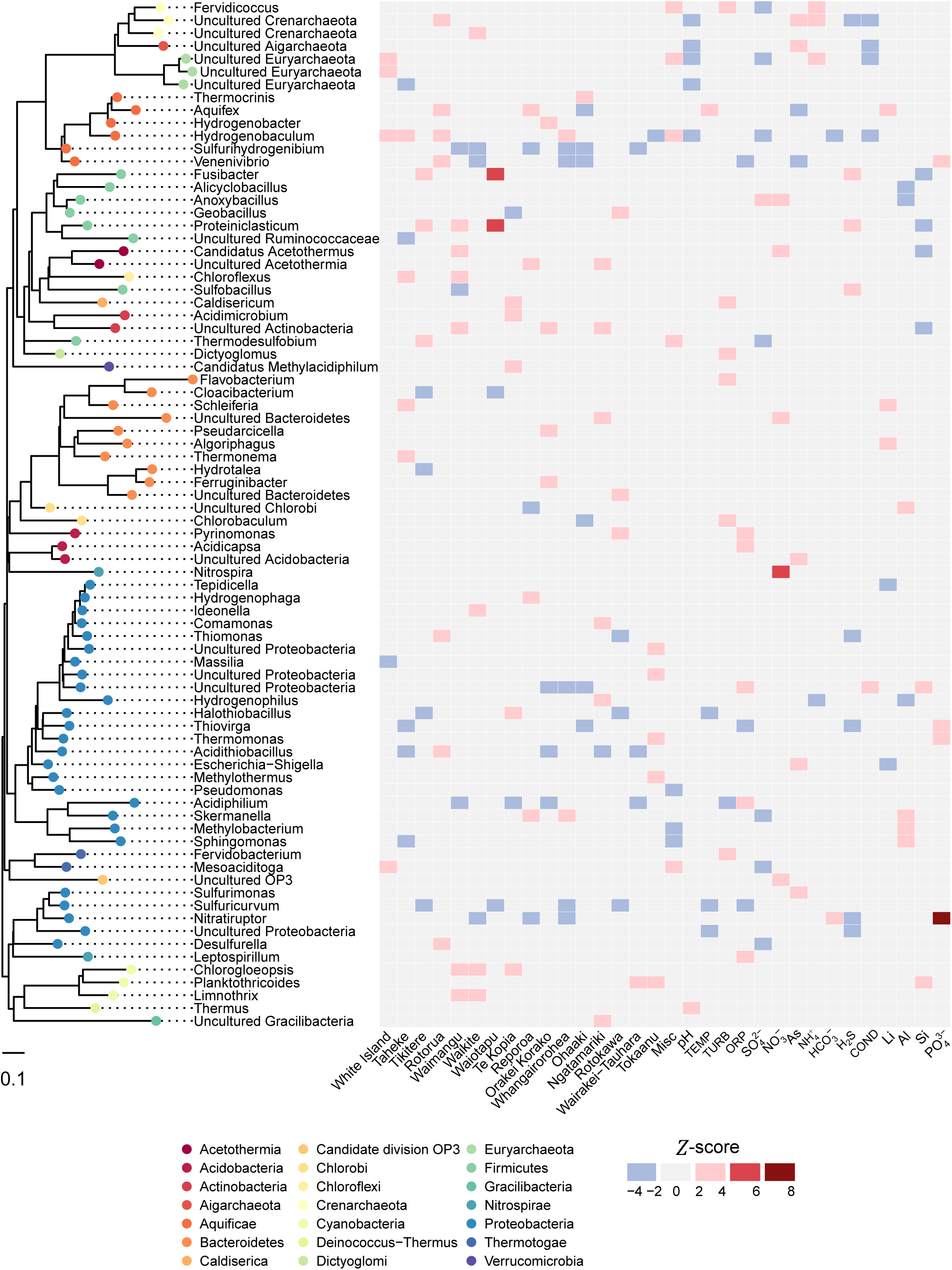
Taxonomic association with location and physicochemsitry. The heat map displays positive (red) and negative (blue) association of genus-level taxa (> 0.1 % average relative abundance) with each geothermal field and significant environmental variables, based on *Z*-scores of abundance log ratios. Each taxon is colour-coded to corresponding phylum on the approximately maximum-likelihood phylogenetic tree.

**Fig. 5.**
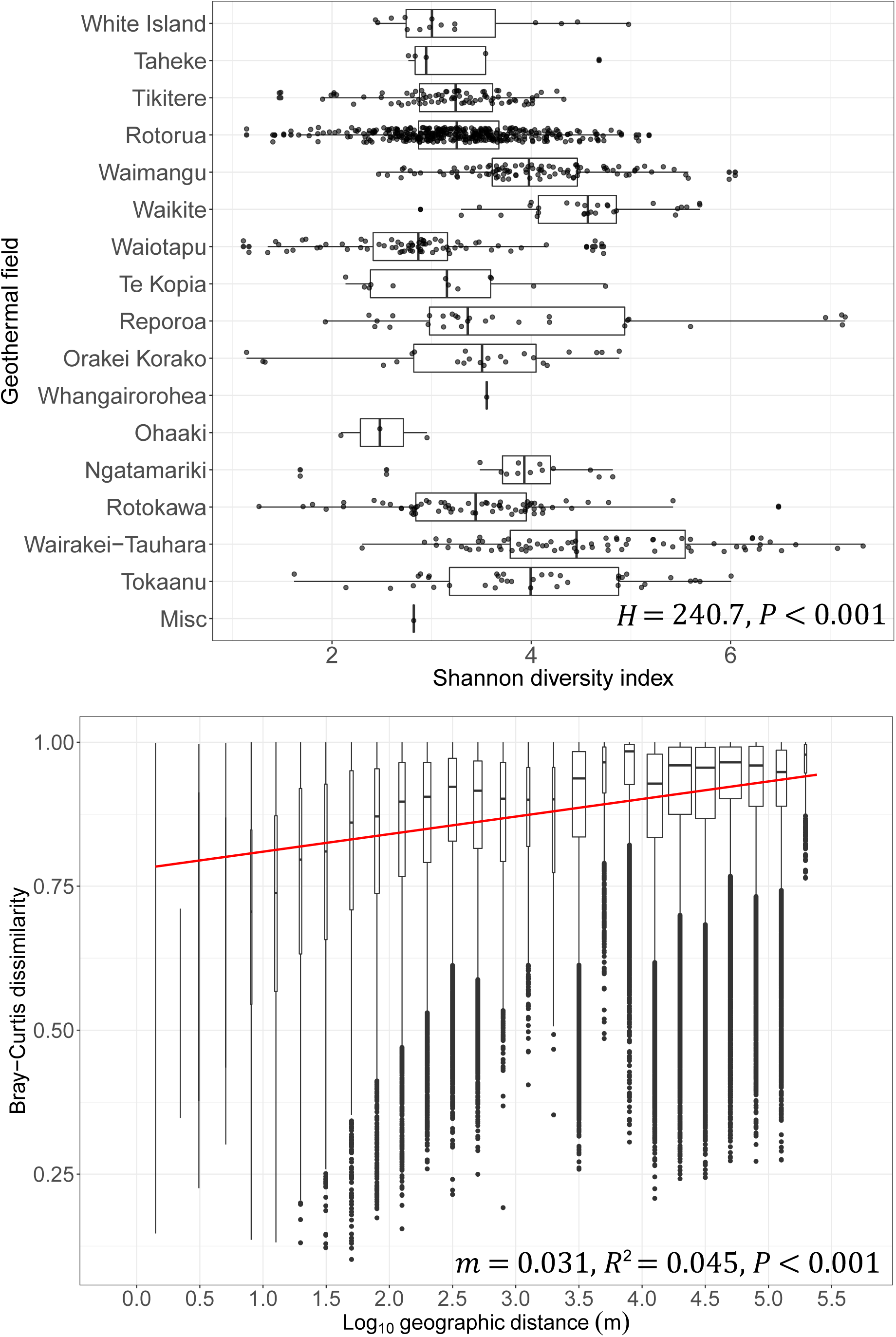
Alpha and beta diversity against geographic distance. (Top) Alpha diversity scales (via Shannon index) across individual springs, separated by geothermal fields. Fields are ordered from north to south (*H*: Kruskal-Wallis test). (Bottom) A distance-decay pattern of beta diversity (via Bray-Curtis dissimilarities of 925 springs) against pairwise geographic distance in metres, with linear regression applied. Geographic distance is split into bins to aid visualisation of the spread.

### Aquificae and Proteobacteria taxa are abundant and widespread

In order to determine whether individual microbial taxa favoured particular environmental conditions and locations, we first assessed the distribution of genera across all individual springs. Within 17 geothermal fields and 925 geothermal features, 21 phyla were detected with an average relative abundance > 0.1 % (Fig. 4). Surprisingly, we found that two phyla and associated genera, Proteobacteria (*Acidthiobacillus* spp.) and Aquificae (*Venenivibrio, Hydrogenobaculum*, *Aquifex* spp.), dominated these ecosystems (65.2 % total average relative abundance across all springs), composing nine of the 15 most abundant genera > 1 % average relative abundance (Table 1). Considering the broad spectrum of geothermal environmental conditions sampled in this study (we assessed microbial communities in springs across a pH gradient of nine orders of magnitude and a temperature range of ~ 87°C), this result was surprising and we believe unprecedented in the literature. Proteobacteria was the most abundant phylum across all samples (34.2 % of total average relative abundance; Table 1), found predominantly at temperatures less than 50 °C (Supplementary Fig. 8). Of the 19 most abundant proteobacterial genera (average relative abundance > 0.1 %), the majority are characterised as aerobic chemolithoautotrophs, utilising either sulfur species and/or hydrogen for metabolism. Accordingly, the most abundant (11.1 %) and prevalent (62.9 %) proteobacterial genus identified was *Acidithiobacillus*, a moderately thermophilic, acidophilic autotroph that utilises reduced sulfur compounds, iron or hydrogen as energy for growth.

**Table 1.** Average relative abundances and prevalence of phyla and genera. Only taxa above a 1 % average compositional threshold are shown. Maximum abundance of each taxon within individual features and standard deviation are noted. Where taxonomy assignment failed to classify to genus level, the closest assigned taxonomy is shown (f = family, o = order, p = phylum).

| Phylum | Genus | Abundance | SD | Max | Prevalence |
| --- | --- | --- | --- | --- | --- |
| Aquificae | *Venenivibrio* | 0.112 | 0.231 | 0.968 | 0.742 |
| Proteobacteria | *Acidithiobacillus* | 0.111 | 0.242 | 0.994 | 0.629 |
| Aquificae | *Hydrogenobaculum* | 0.100 | 0.235 | 0.999 | 0.608 |
| Aquificae | *Aquifex* | 0.086 | 0.212 | 0.971 | 0.497 |
| Deinococcus-Thermus | *Thermus* | 0.025 | 0.071 | 0.732 | 0.552 |
| Proteobacteria | *Thiomonas* | 0.024 | 0.101 | 0.941 | 0.396 |
| Proteobacteria | *Desulfurella* | 0.022 | 0.067 | 0.758 | 0.497 |
| Crenarchaeota | *Sulfolobaceae* (f) | 0.020 | 0.091 | 0.951 | 0.416 |
| Euryarchaeota | *Thermoplasmatales* (o) | 0.019 | 0.059 | 0.495 | 0.539 |
| Proteobacteria | *Thiovirga* | 0.015 | 0.077 | 0.816 | 0.374 |
| Proteobacteria | *Hydrogenophilaceae* (f) | 0.015 | 0.072 | 0.704 | 0.406 |
| Thermodesulfobacteria | *Caldimicrobium* | 0.015 | 0.052 | 0.651 | 0.519 |
| Proteobacteria | *Hydrogenophilus* | 0.013 | 0.045 | 0.432 | 0.484 |
| Thermotogae | *Mesoaciditoga* | 0.011 | 0.033 | 0.286 | 0.410 |
| Parcubacteria | *Parcubacteria* (p) | 0.010 | 0.024 | 0.193 | 0.608 |

Aquificae (order Aquificales) was the second most abundant phylum overall (31 % average relative abundance across 925 springs) and included three of the four most abundant genera; *Venenivibrio*, *Hydrogenobaculum* and *Aquifex* (11.2 %, 10.0 % and 8.6 % respectively; Table 1). As the Aquificae are thermophilic (T_opt_ 65 – 85 °C)^50^, they were much more abundant in warmer springs (> 50 °C; Supplementary Fig. 8). The minimal growth temperature reported for characterised Aquificales species (*Sulfurihydrogenibium subterraneum* and *S. kristjanssonii*)^50^ is 40 °C and may explain the low Aquificae abundance found in springs less than 50 °C. Terrestrial Aquificae are predominately microaerophilic chemolithoautotrophs that oxidise hydrogen or reduced sulfur compounds; heterotrophy is also observed in a few representatives^50^. Of the 14 currently described genera within the Aquificae, six genera were relatively abundant in our dataset (average relative abundance > 0.1 %; Fig. 4): *Aquifex*, *Hydrogenobacter*, *Hydrogenobaculum* and *Thermocrinis* (family Aquificaceae); and *Sulfurihydrogenibium* and *Venenivibrio* (family Hydrogenothermaceae). No signatures of the Desulfurobacteriaceae were detected. This is consistent with reports that all current representatives from this family are associated with deep-sea or coastal thermal vents^50^. *Venenivibrio* (OTUs; *n* = 111) was also the most prevalent and abundant genus across all communities (Table 1). This taxon, found in 74.2 % (*n* = 686) of individual springs sampled, has only one cultured representative, *V. stagnispumantis* (CP.B2^T^), which was isolated from the Waiotapu geothermal field in the TVZ^31^. The broad distribution of this genus across such a large number of habitats was surprising, as growth of the type strain is only supported by a narrow set of conditions (pH 4.8 – 5.8, 45 – 75 °C). Considering this and the number of *Venenivibrio* OTUs detected, we interpret this result as evidence there is substantial undiscovered phylogenetic and physiological diversity within the genus. The ubiquity of *Venenivibrio* suggests that either the metabolic capabilities of this genus extend substantially beyond those described for the type strain, and/or that many of the divergent taxa could be persisting and not growing under conditions detected in this study^51,52^.

### Fine-scale geochemical and geographical associations exist at the genus level

The two most abundant phyla, Proteobacteria and Aquificae, were found to occupy a characteristic ecological niche (< 50 °C and > 50 °C respectively, Supplementary Fig. 8). To investigate specific taxa-geochemical associations beyond just temperature and pH, we applied a linear model to determine enrichment of taxa in association with geothermal fields and other environmental data (Fig. 4). The strongest associations between taxa and chemistry (*Z*-score > 4) were between *Nitrospira*–nitrate 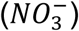 and *Nitratiruptor*–phosphate 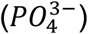. *Nitrospira* oxidises nitrite to nitrate and therefore differential high abundance of this taxon in nitrate-rich environments is expected. Further, the positive *Nitratiruptor*–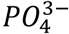 relationship suggests phosphate is a preferred nutritional requirement for this chemolithoautotroph^53^ and informs future efforts to isolate members of this genus would benefit from additional phosphate or the presence of reduced P compounds in the culture medium^54,55^. *Thermus* and *Hydrogenobaculum* were the only bacterial taxa to differentially associate (compared to other taxa) positively and negatively with pH respectively. This is consistent with the lack of acidophily phenotype (pH < 4) reported in *Thermus* spp.^56^ and the preferred acidic ecological niche of *Hydrogenobaculum*^57^. *Aquifex* was the only genus to display above average association with temperature, confirming abundance of this genera is significantly enhanced by hyperthermophily^58^.

Similar to the chemical-taxa associations discussed above, differential abundance relationships were calculated with respect to individual geothermal fields (Fig. 4). The Rotorua geothermal field, which contains springs across the pH scale, was closely associated with the highly abundant and prevalent *Acidithiobacillus* and *Venenivibrio*. On the other hand, Te Kopia, a predominantly acidic geothermal system, produced the only positive associations with *“Methylacidiphilum”* (Verrucomicrobia), *Acidimicrobium* (Actinobacteria), *Terrimonas* (Bacteroidetes) and *Halothiobacillus* (Proteobacteria). Curiously, the strongest positive taxa-geography associations were identified between both *Fusibacter*-Waiotapu and *Proteiniclasticum*-Waiotapu. Given the Waiotapu geothermal field is predominately an acid-sulfate system, the association of Waiotapu to these anaerobic, mesophilic neutrophiles was unexpected, although a species of *Fusibacter* has been isolated from a mesophilic spring^59,60^. These relationships are likely describing sub-community requirements that are otherwise not captured by conventional spatial-statistical analysis, therefore providing insight into previously unrecognised microbe-niche interactions.

### Microbial distance-decay patterns differ at local and regional scales

Environmental selection, ecological drift, diversification and dispersal limitation all contribute to distance-decay patterns^4,61^. While several recent studies have shown microbial dispersal limitations and distance-decay patterns exist in diverse environments^9,21,61,62^, the point of inflection between dispersal limitation and selection, at regional and local geographic scales, remains under-studied. We identified a positive distance-decay trend with increasing geographic distance between 925 geothermal spring communities across the TVZ region (*m* = 0.031, *P* < 0.001; Fig. 5). This finding strongly suggests dispersal limitation exists between individual geothermal fields. Increasing the resolution to within individual fields, distance-decay patterns are negligible compared to the regional scale (Supplementary Table 6). Interestingly, the greatest pairwise difference (y = 1) between Bray-Curtis dissimilarities was also observed in springs classified as geographically-adjacent (< 1.4 m). In the 293 springs pairs separated by < 1.4 m, temperature had a greater correlation with beta diversity than pH (Spearman’s coefficient: *ρ* = 0.44 and 0.30 respectively, *P* < 0.001). This result illustrates the stark spatial heterogeneity and selective processes that can exist within individual geothermal fields. Congruently, each OTU was detected in an average of only 13 springs (Supplementary Fig. 4). We propose that physical dispersal within geothermal fields is therefore not limiting, but the physicochemical diversity of hotsprings acts as a barrier to the colonisation of immigrating taxa. However, even between some neighbouring springs with similar (95% *CI*) geochemical signatures, we did note some dissimilar communities were observed (for example, Waimangu geothermal field; Fig. 3E). These differing observations can be explained either one of two ways; firstly, the defining parameter driving community structure was not one of the 46 physicochemical variables measured in this study (e.g. dissolved organic carbon); or secondly, through the process of dispersal, the differential viability of some extremophilic taxa restricts gene flow and contributes to population genetic drift within geothermal fields^63,64^. We often consider “extremophilic” microorganisms living in these geothermal environments as the epitome of hardy and robust. In doing so, we overlook that their proximal surroundings (i.e. immediately outside the host spring) may not be conducive to growth and survival^65^ and therefore the divergence of populations in neighbouring, chemically-homogenous spring ecosystems is plausible. Future work could include understanding individual population response^66^ to these community-wide selective pressures.

## Conclusion

This study presents data on both niche and neutral drivers of microbial biogeography in 1,000 geothermal springs at a near-national scale. Our comprehensive data set, with sufficient sampling density and standardised methodology, is the first of its kind to enable a robust spatio-chemical statistical analysis of microbial communities at the regional level across broad physicochemical gradients. Unequivocally, pH drives diversity and community complexity structures within geothermal springs. This effect, however, was only significant at temperatures < 70 °C. We also identified specific taxa associations and finally demonstrated that geochemical signatures can be indicative of community composition. Although a distance-decay pattern across the entire geographic region indicated dispersal limitation, the finding that 293 adjacent community pairs exhibited up to 100 % dissimilarity suggests niche selection drives microbial community composition at a localised scale (e.g. within geothermal fields). This research provides a comprehensive dataset that should be used as a foundation for future studies (e.g. diversification^66^ and drift^67,68^ elucidation on targeted spring taxa). It complements the recently published Earth Microbiome Project^45^ by expanding our knowledge of the biogeographical constraints on aquatic ecosystems using standardised quantification of broad physicochemical spectrums. There is also potential to use the two studies to compare geothermal ecosystems on a global scale. Finally, our research provides a springboard to assess the cultural, recreational and resource development value of the microbial component of geothermal springs, both in New Zealand and globally. Many of the features included in this study occur on culturally-important and protected land for Māori, therefore this or follow-on future projects may provide an avenue for exploration of indigenous knowledge, while assisting in conservation efforts and/or development.

## Methods

### Field sampling & processing

Between July 2013 and April 2015, 1,019 aqueous samples were collected from 974 distinct geothermal features within 18 geothermal fields in the TVZ. A three litre integrated water column sample was taken from each geothermal spring, lake, stream, or the catchment pool of geysers for microbial and chemical analyses. Comprehensive physical and chemical measurements, and field observational metadata were recorded contemporaneously with a custom-built application and automatically uploaded to a database. All samples were filtered within two hours of collection and stored accordingly (Supplementary Table 7). Total DNA was extracted using a modified CTAB method^69^ with the PowerMag Microbial DNA Isolation Kit using SwiftMag technology (MoBio Laboratories, Carlsbad, CA, USA). The V4 region of the 16S rRNA gene was amplified in triplicate using universal Earth Microbiome Project^70^ primers F515 (5’-GTGCCAGCMGCCGCGGTAA-3’) and R806 (5’-GGACTACVSGGGTATCTAAT-3’). SPRIselect (Beckman Coulter, Brea, CA, USA) was used to purify DNA following amplification. Amplicon sequencing was performed using the Ion PGM System for Next-Generation Sequencing with the Ion 318v2 Chip and Ion PGM Sequencing 400 Kits (ThermoFisher Scientific, Waltham, MA, USA).

Forty-seven separate physicochemical parameters were determined for each hotspring sample collected. Inductively coupled plasma-optical emission spectrometry (ICP-OES) and -mass spectrometry (ICP-MS) were used to determine the concentrations of aqueous metals and non-metals (31 species), and various UV-Vis spectrometry methods were used to determine aqueous nitrogen species 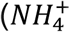, 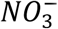, 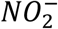, 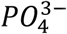, with *Fe^2+^*, *H_2_*, 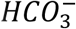 and *Cl*^−^ determined via titration, and sulfate concentration measured via ion chromatography (IC). Conductivity (COND), dissolved oxygen (dO), oxidation-reduction potential (ORP), pH, temperature (TEMP), and turbidity (TURB) were determined using a Hanna Instruments (Woonsocket, RI, USA) multiparameter field meter *in situ*. Expanded details on sampling procedures, sample processing, DNA extraction, DNA amplification, and chemical analyses can be found in the Supplementary Methodology and Supplementary Table 7.

### DNA sequence processing

DNA sequences were processed through a custom pipeline utilising components of UPARSE^71^ and QIIME^72^. An initial screening step was performed in mothur^73^ to remove abnormally short (< 275 bp) and long (> 345 bp) sequences. Sequences with long homopolymers (> 6) were also removed. A total of 47,103,077 reads were quality filtered using USEARCH v7^71^ with a maximum expected error of 1 % (fastq_maxee = 2.5) and truncated from the forward primer to 250 bp. Retained sequences (85.4 % of initial reads) were dereplicated and non-unique sequences removed. Next, reads were clustered to 97 % similarity and chimera checked using the cluster_otus command in USEARCH, and a *de novo* database was created of representative operational taxonomic units (OTUs). 93.2 % of the original pre-filtered sequences (truncated to 250 bp) mapped to these OTUs, and taxonomy was assigned using the Ribosomal Database Project Classifier^74^ (with a minimum confidence score of 0.5) against the SILVA 16S rRNA database (123 release, July 2015)^75^. The final read count was 43,202,089, with a mean of 43,905 reads per sample. Chloroplasts and mitochondrial reads were removed (1.0 and 0.5 % respectively of the final read count) and rarefaction was performed to 9,500 reads per sample.

### Statistical analyses

All statistical analyses and visualisation were performed in the R environment^76^ using phyloseq^77^, vegan^78^ and ggplot2^79^ packages. Alpha diversity was calculated using the estimate_richness function in phyloseq. A series of filtering criteria were applied to the 46 geochemical parameters measured in this study to identify metadata that significantly correlated with alpha diversity in these spring communities. First, collinear variables (Pearson correlation coefficient | *r* | > 0.7) were detected^80^. The best-fit linear regression between alpha diversity (using Shannon’s index) and each variable was used to pick a representative from each collinear group. This removed variables associating with the same effect in diversity. Multiple linear regression was then applied to remaining variables, before and after a stepwise Akaike information criterion (AIC) model selection was run^81^. Due to the wide pH, temperature and geographic ranges for this dataset, samples were also binned by increments of each criterion respectively (Supplementary Fig. 1), with non-parametric Kruskal-Wallis (*H*) testing performed to identify any significant differences between groups. Finally, correlation of pH and temperature against Shannon diversity was calculated using Pearson’s coefficient | *r* |.

Bray-Curtis dissimilarity was used for all beta diversity comparisons. For ordination visualisations, a square-root transformation was applied to OTU relative abundances prior to non-metric multidimensional scaling (k = 2) using the metaMDS function in the vegan package. ANOSIM (|*R*|) was used to compare beta diversity across the same pH, temperature and geographic groups (i.e. geothermal fields) used for alpha diversity analyses, followed by pairwise Wilcox testing with Bonferroni correction to highlight significance between individual groups. Linear regression was applied to pairwise geographic distances against spring community dissimilarities to assess the significance of distance-decay patterns. These comparisons were similarly performed on spring communities constrained to each geothermal field. A second series of filtering criteria was applied to geochemical parameters to identify metadata that significantly correlated with beta diversity. Mantel tests were performed between beta diversity and all 46 physicochemical variables using Spearman’s correlation coefficient (*ρ*) with 9,999 permutations. In decreasing order of correlation, metadata were added to a PERMANOVA analysis using the adonis function in vegan.

Metadata significantly correlating with beta diversity (*P* < 0.01) was assessed for collinearity using Pearson’s coefficient | *r* |^80^. In each collinear group (|*r* | > 0.7), the variable with the highest mantel statistic was chosen as the representative. Low variant geochemical variables (*σ* < 0.25 ppm) were then removed to allow a tractable number of explanatory variables for subsequent modelling. Constrained correspondence analysis (using the cca function in vegan) was then applied to OTUs, geothermal field locations and the reduced set of metadata. OTUs were first agglomerated to their respective genera (using the tax_glom function in phyloseq) and then low abundant taxa (< 0.7 %) of total mean taxon abundance were removed. Typical geochemical signatures within each geothermal field were used to produce ternary diagrams of *Cl^−^*, 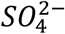 and 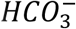 ratios using the ggtern package^82^.

Finally, to detect significant associations between taxa, geochemistry and other metadata (i.e. geothermal field observations), a linear model was applied to determine log enrichment of taxa using edgeR^83^. To simplify the display of taxonomy in this model, we first agglomerated all OTUs to their respective genera or closest assigned taxonomy group (using the tax_glom function in phyloseq), and then only used taxa present in at least 5 % of samples and > 0.1 % average relative abundance. Log fold enrichments of taxa were transformed into *Z*-scores and retained if absolute values were > 1.96. Results were visualized using ggtree^84^. A phylogenetic tree was generated in QIIME by confirming alignment of representative OTU sequences using PyNAST^85^, filtering the alignment to remove positions which were gaps in every sequence and then building an approximately maximum-likelihood tree using FastTree^86^ with a midpoint root.

## Data availability

Raw sequences have been deposited into the European Nucleotide Archive (ENA) under study accession number PRJEB24353. General data is presented in a user-friendly queryable website (http://1000springs.org.nz). All code used for statistics and figures is available through GitLab (https://gitlab.com/morganlab/collaboration-1000Springs/1000Springs).

## Acknowledgements

The authors wish to acknowledge all our landowners and Māori collaborators for access and support of this research. *Mana whenua* (customary rights) is acknowledged for all data generated arising from geothermal features within *rohe* of iwi. The primary collection and processing of samples was funded by an MBIE Smart Idea grant (C05X1203 - Microbial Bioinventory of Geothermal Ecosystems) awarded to MBS and SCC, colloquially known as the 1,000 Springs Project (http://1000springs.org.nz). JFP was also supported by a GNS Science Postgraduate Scholarship (under the Geothermal Resources of New Zealand Research Programme), and the University of Waikato Hilary Jolly Memorial Scholarship. We thank Karen Houghton and Hanna-Annette Peach for assistance with field work, Kevin Lee for advice on bioinformatics, and Jayadev Payyakkal Viswam for help with DNA extractions. We also thank Isabelle Chambefort, Ed Mroczek and Nellie Olsen for valuable comments.

## Author contributions

MBS, SCC, JFP, IRM and MDC designed the study. JFP, DWE, MBS, CRC and GLJW undertook field work and processing of samples. MB, DW, MBS, JFP and AMH designed the field application, database and website. GLJW performed DNA extractions and sequencing. JFP and CKL processed DNA sequences. JFP and XCM performed data analysis and statistics. JFP, CRC and MBS wrote the manuscript, with assistance from SCC, XCM, CKL, IRM and GLJW.

